# Addressing persistent challenges in digital image analysis of cancerous tissues

**DOI:** 10.1101/2023.07.21.548450

**Authors:** Sandhya Prabhakaran, Clarence Yapp, Gregory J. Baker, Johanna Beyer, Young Hwan Chang, Allison L. Creason, Robert Krueger, Jeremy Muhlich, Nathan Heath Patterson, Kevin Sidak, Damir Sudar, Adam J. Taylor, Luke Ternes, Jakob Troidl, Yubin Xie, Artem Sokolov, Darren R. Tyson, the Cell Imaging Hackathon 2022 Participants (Alphabetical authorship, apart from primary contributors)

**Affiliations:** H. Lee Moffitt Cancer Center and Research Institute; Laboratory of Systems Pharmacology, Harvard Medical School; Harvard Medical School; School of Engineering and Applied Sciences, Harvard University; Computational Biology Program, Department of Biomedical Engineering, Oregon Health & Science University, Portland; Knight Cancer Institute, Department of Biomedical Engineering, Oregon Health & Science University, Portland; Harvard University; Aspect Analytics; Quantitative Imaging Systems; Sage Bionetworks; Memorial Sloan Kettering Cancer Center; Department of Pharmacology, Vanderbilt University, Nashville, TN

**Keywords:** Multiplexed images, cancer, image analysis, domain representation, artifact removal, scalability, thumbnail generation

## Abstract

The National Cancer Institute (NCI) supports many research programs and consortia, many of which use imaging as a major modality for characterizing cancerous tissue. A trans-consortia Image Analysis Working Group (IAWG) was established in 2019 with a mission to disseminate imaging-related work and foster collaborations. In 2022, the IAWG held a virtual hackathon focused on addressing challenges of analyzing high dimensional datasets from fixed cancerous tissues. Standard image processing techniques have automated feature extraction, but the next generation of imaging data requires more advanced methods to fully utilize the available information. In this perspective, we discuss current limitations of the automated analysis of multiplexed tissue images, the first steps toward deeper understanding of these limitations, what possible solutions have been developed, any new or refined approaches that were developed during the Image Analysis Hackathon 2022, and where further effort is required. The outstanding problems addressed in the hackathon fell into three main themes: 1) challenges to cell type classification and assessment, 2) translation and visual representation of spatial aspects of high dimensional data, and 3) scaling digital image analyses to large (multi-TB) datasets. We describe the rationale for each specific challenge and the progress made toward addressing it during the hackathon. We also suggest areas that would benefit from more focus and offer insight into broader challenges that the community will need to address as new technologies are developed and integrated into the broad range of image-based modalities and analytical resources already in use within the cancer research community.

## Introduction

The cellular histopathology of cancer has been studied for hundreds of years with the first description of a cellular origin of cancer being described by Rudolph Virchow in the 1850s^1^; histopathologic evaluation of tumor samples is now an essential component of standard clinical practice. The analysis of cancerous tissue samples has long been performed with direct visual inspection by trained pathologists^2^, but, with the technological advancement of instruments to spatially resolve tissue samples and the concomitant increase in computational power and function, a new field of computational image analysis of cancerous tissues has evolved. With this explosion in our ability to generate digital images of cancer has come the challenge of how to quantitatively extract meaningful features from them in an automated way.

The National Cancer Institute (NCI) broadly supports research programs that foster emerging areas in cancer biology and the development of new experimental models for cancer research, including the Cancer Systems Biology Consortium (CSBC), Physical Sciences–Oncology Network (PS–ON), the Human Tumor Atlas Network (HTAN) and the more recent additions of the Cellular Cancer Biology Imaging Research (CCBIR), Acquired Resistance to Therapy Network (ARTNet), Patient-Derived Xenograft Network (PDXNet), and TaRget Enablement to Accelerate Therapy Development for AD (TREAT-AD). Imaging is a core technology among many research groups within these programs. In addition, non-cancer-specific initiatives such as Human BioMolecular Atlas Project (HuBMAP)^3,4^ and the Human Cell Atlas (HCA)^5–7^ rely on digital imaging as a major component of their programs and are generating enormous amounts of spatially resolved data with up to hundreds or thousands of measurements mappable to individual cells, at an ever-increasing scale. Standard image processing techniques of image alignment and stitching, segmentation and cell type calling have facilitated automated extraction of quantitative features from digital images^8–11^, and their further refinement will continue extending their utility and capabilities. However, the next generation of imaging data comprises many more measurements per cell and covers an ever-expanding volume of tissue per sample. The interpretation of imaging data may therefore be limited when analyzed using the traditional image processing pipeline that does not utilize the richer features of modern imaging data.

In response to the changing landscape of digital image analysis across NIH-funded programs, a trans-consortia Image Analysis Working Group (IAWG) was initiated in 2019 with the objectives of disseminating the work of the multitude of imaging-related endeavors across all of the research groups and developing collaborations to address common challenges. In January 2020, the IAWG held a workshop designed to identify challenges that could be addressed within an in-person hackathon setting, which was hosted by Vanderbilt University in early March, 2020, a few days prior to the nationwide shutdown in response to COVID-19. Within the proceedings of the IAWG’s combined workshop/hackathon we described several specific challenges centered around the traditional image processing pipeline and how they were addressed by the hackathon^12^, and the IAWG has continued to identify and address outstanding challenges in the digital image analysis pipeline.

In this perspective, we discuss what we view as important open questions in the analysis of multiplexed tissue images, the first steps toward deeper understanding of these questions, what possible solutions may look like, what approaches were addressed during the Image Analysis Hackathon 2022, and where further work remains. The hackathon considered eleven challenges that fell into three main themes: 1) challenges to cell type classification and assessment, 2) translation and visual representation of spatial aspects of high dimensional data, and 3) scaling digital image analyses to large (multi-TB) datasets. We discuss our efforts toward addressing specific aspects of these main themes and attempt to provide a broader view of the remaining challenges to a more robust automated pipeline for large-scale multiplexed digital image analysis.

### Challenges to identifying and classifying cell types

Light microscopy-based digital histopathological analysis of cancer tissues has been greatly enhanced by several recently-developed multiplexed techniques, including antibody-based methods such as cyclic immunofluorescence (CyCIF)^13^, iterative indirect immunofluorescence imaging (4i)^14^, imaging mass spectrometry (iMS), IMC (Imaging Mass Cytometry)^15^, multiplexed immunofluorescence (MxIF), and co-detection by indexing^16^ (CODEX); and spatial transcriptomic techniques such as multiplexed error-robust fluorescence *in situ* hybridization (MERFISH)^17^ and sequential fluorescence *in situ* hybridization (seqFISH)^18^. These techniques increase the number of features captured per cell and have greatly expanded our understanding of the high level of variation that exists among cells, including those with very similar morphological characteristics. However, accurate quantification of these features and their association with individual cells within the spatial context of cancerous tissue largely relies on image segmentation and other techniques used to demarcate cellular boundaries. While deep learning approaches have been successfully deployed to automate this task with previously unattainable levels of accuracy (*e.g.*, CellPose^9^, Mesmer^19^, Hover-Net^20^), numerous challenges persist that limit our ability to automatically achieve accurate quantification of features at single-cell and sub-cellular resolution. Such limitations include tissue handling artifacts (e.g., bubbles, dust and debris, crush and sectioning artifacts, etc.); technical artifacts (e.g., inaccurate image stitching, uneven illumination, and antibody aggregates) (**Figure 1A**); and image projection artifacts that arise while projecting imprecise and overlapping cell boundaries from three-dimensional structures to two dimensions resulting in apparent “lateral spillover^21^.” All of these can contribute to errors in the output of segmentation, gating, and cell type calling algorithms that generally expect ideal data. Moreover, these high-dimensional feature sets must be analyzed and visualized to compare the differences and similarities among cell types, yet no well-accepted standardized method for these comparisons (or how to define each cell type) has been described. Within the hackathon setting, several teams worked to specifically address some aspects of these issues by developing new or leveraging existing tools to suppress the contribution of these artifacts to the process of cell type classification and visualization of the resulting classes. The specific objectives of these teams were 1) to develop a process for automatically detecting image artifacts, with either a human-in-the-loop or a completely automated process; 2) to suppress the effects of image artifacts on downstream analyses; 3) to correct the effects of lateral spillover on cell type classification; and 4) to optimize the automated visualization of high dimensional single-cell data representative of distinct cell classes.

**Figure 1:**
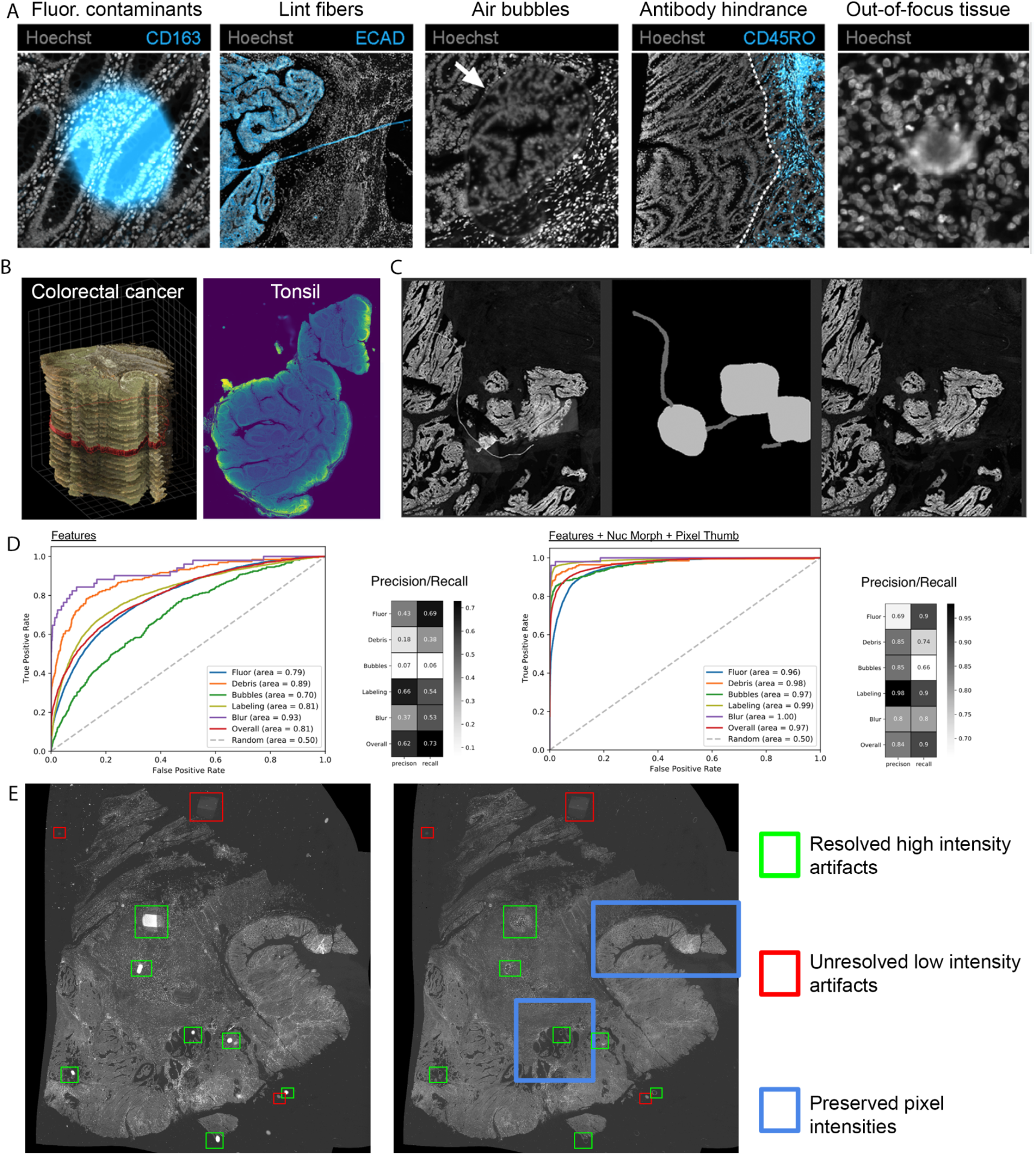
Strategies for artifact detection and correction. **A**) Examples of common imaging artifacts in fluorescence microscopy. From left to right: miscellaneous fluorescent contaminants, autofluorescent lint fibers, air bubbles causing refractive index mismatch, antibody hindrance (broad region of low antibody reactivity), and out-of-focus tissue. **B**) CyCIF datasets used for the artifact-related hackathon challenges, featuring human colorectal cancer and tonsil tissue. **C**) A fibrous artifact and illumination errors are visible (*left*) and manually annotated (*middle*) to facilitate its detection and suppression (*right*). **D**) ROC curve analysis for artifact detection performance of a multilayer perceptron trained on mean immunomarker signals alone (Features, FS1 in main text, *left)*, or Features plus segmentation-based nuclear morphology attributes (Nuc Morph) and pixel-level image statistics (Pixel Thumb; FS3 in main text, *right*). Also see Supplementary Figure 2. **E**) Comparison of before (*left*) and after (*right*) automatic artifact correction. Artifacts that have been sufficiently removed or unresolved are shown with green or red boxes respectively. Image intensity was left unperturbed by artifact correction in tissues without artifacts. Examples shown with blue boxes.

The participants were provided with CyCIF datasets representing two tissue types - colorectal cancer (Figure 1B, *left*) and tonsil (Figure 1B, *right*) acquired at 0.65 micrometers per pixel with a 20X/0.75NA objective lens. The colorectal cancer data was acquired as part of the HTAN efforts and consisted of 25 serial sections stained with 10 rounds of a 30-plex CyCIF^22^. The tonsil dataset underwent 9 rounds of CyCIF and was used in the development of the Multiple-choice microscopy pipeline^10^ (MCMICRO). These datasets provide examples of the variety of imaging artifacts that plague samples even under the best conditions, providing a suitable resource for the development of tools to address the challenges listed above.

### Automatic artifact detection from the spatial feature table

Experimental artifacts alter the natural dynamic range of signal intensities and cause false positive signals in derived single-cell data (Figure 1A). An ideal quality control (QC) tool for quantitative multiplex microscopy should automatically account for artifacts imposed at all stages in the data acquisition pipeline, from pre-analytical variables such as biospecimen quality and tissue fixation conditions, to errors in tile stitching, imaging alignment, and cell segmentation. However, the majority of QC tools for digital pathology are limited to strategies for evaluating a subset of artifacts induced by sample preparation or data acquisition such as out-of-focus imaging, tissue degradation, and batch effect^23–28^. They fail to account for downstream errors in image processing such as tile stitching, image registration, and cell segmentation. One such tool for identifying and removing artifacts in tissue-derived single-cell data from sample preparation to image acquisition and processing is CyLinter^29^, a tool designed for human-in-the-loop artifact curation. While manual artifact curation is effective for limited numbers of samples (<20) probed with relatively few immunomarkers (<20), it tends to scale poorly with dataset size. Thus, methods for automated detection of microscopy artifacts are needed to enhance workflow efficiency, minimize curator burden, and mitigate human bias.

This challenge asked participants to use machine learning methods to automatically detect visual artifacts in multiplex images (Supplementary Figure 1). Multiple supervised modeling approaches were assessed for their ability to automatically identify each of five different artifact categories routinely encountered in multiplex immunofluorescence (IF) images: 1) fluorescent contaminants, 2) uneven immunolabeling, 3) coverslip air bubbles, 4) slide debris, and 5) image blur (Figure 1A). Ground truth annotations for the different artifact categories were generated through manual curation and provided to challenge participants, allowing them to establish three different feature sets (FS) for model training. The first comprised per-cell mean signal intensities from the colorectal cancer image, spanning 21 immunomarker channels plus Hoechst nuclear dye (FS1). The second combined the features in FS1 with an additional 8 nuclear morphology attributes derived from cell segmentation outlines (FS2), and the third consisted of a combination of FS1 and FS2 plus 289 pixel-level summary statistics calculated on 30-pixel X 30-pixel thumbnail multi-channel images cropped from the full image (FS3).

Participants began by scaling the feature data using standard approaches and evaluating decision-boundary classifiers such as linear and quadratic discriminant analysis (LDA and QDA), partial least squares-discriminant analysis (PLS-DA)^30^, and support vector machines (SVM)^31^, but these models were found to have poor predictive power when compared to ensemble models such as random forests (RFs)^32^, multi-layer perceptrons (MLPs)^17^, and boosted trees^33^ (Figure 1D; Supplementary Figure 2). Notably, the addition of nuclear morphology features (FS2) did not significantly improve any of the algorithms compared to the mean intensity feature table (FS1), whereas the addition of pixel-level features (FS3) resulted in a dramatic improvement in accuracy (Figure 1D; Supplementary Figure 2).

### Artifact correction/suppression directly on images

Visual examination of digital images is often used to assess spatial patterns at cellular and subcellular resolution. The artifact detection and removal using annotations described above does not alter the images themselves but rather the extracted information, and since artifacts generally appear as large/abundant high intensity objects, they can be distracting and lead to incorrect conclusions especially for audiences who are unfamiliar with biological samples or even fluorescence microscopy itself. Thus, correcting or suppressing these artifacts directly in the images would be useful, even if the artifact-induced corruption of the underlying image data cannot be properly retrieved. To this end, different approaches to suppress artifacts were developed and assessed on ground truth annotations (Figure 1C), including deep learning methods that ranged from image in-painting using a pre-trained model to generative adversarial networks (GAN)^34^. The test data consisted of four neighboring registered serial section images with similar content that had not been used in any training sets. Accuracy was based on mean square error (MSE) and peak-signal-to-noise-ratio (PSNR) between suppressed artifacts and a region free from artifacts from a neighboring serial section. The accuracy for the deep learning based methods was low-to-moderate due to the limited training data, although image in-painting produced the best results from the supervised methods. Ultimately, a simpler and more accurate unsupervised approach was found that replaced artifact-corrupted pixels with content from a neighboring section (Figure 1E). Thus, in the absence of artifact annotations, it was still possible to detect artifacts by identifying outliers of intensity across the serial sections, which resulted in comparable accuracy to the deep learning image in-painting approach without requiring huge investment in time and labor to curate training data.

### Lateral spillover correction using REDSEA

Attribution of pixel intensities to different cells in digital images is highly dependent on segmentation accuracy, especially the location of a boundary between a cell and its neighbors (**Figure 2A**). Any errors along this boundary can result in spatial crosstalk of marker signals and lead to nonsensical cell types. The objective for the hackathon was to correct for lateral spillover in a publicly available dataset (CyCIF-processed tonsil sections segmented with Deepcell^35^), assess the quality of the correction, and scale the method to larger image sizes. The approach taken by the participants leveraged a previously devised method called REinforcement Dynamic Spillover EliminAtion (REDSEA)^21^. REDSEA first computes the proportion of the shared boundary between a cell and its neighbors. It then compensates for signal intensity of each channel along that boundary based on the overall expression of that channel in the cell relative to its neighbors. The method was evaluated using sets of cell type markers that are known to be mutually exclusive. Due to the restricted time available during the hackathon, evaluation proceeded using five tiles each for two subsets of the tonsil dataset: 200 × 200 or 800 × 800 pixels. The original implementation of REDSEA was developed using high density tissue and was discovered to crash (with a divide by zero error) when the code encountered isolated cells (i.e., cells with no immediately adjacent cells) in the image. Most isolated cells occurred around the periphery of the image, and is either due to an artifact of cell segmentation or image tiling. Hackathon participants enabled REDSEA to identify these isolated cells as a distinct cluster in high dimensional feature space, while removing them from consideration of lateral spillover (**Figure 2B**). Co-expression plots for pairs of mutually exclusive markers both before and after the modified implementation of REDSEA (**Supplementary Figure 3**) show that we were able to reproduce REDSEA’s results while also increasing the proportion of single-positive cells. This updated implementation has also been translated from MATLAB to Python^36^ to facilitate broader use.

**Figure 2:**
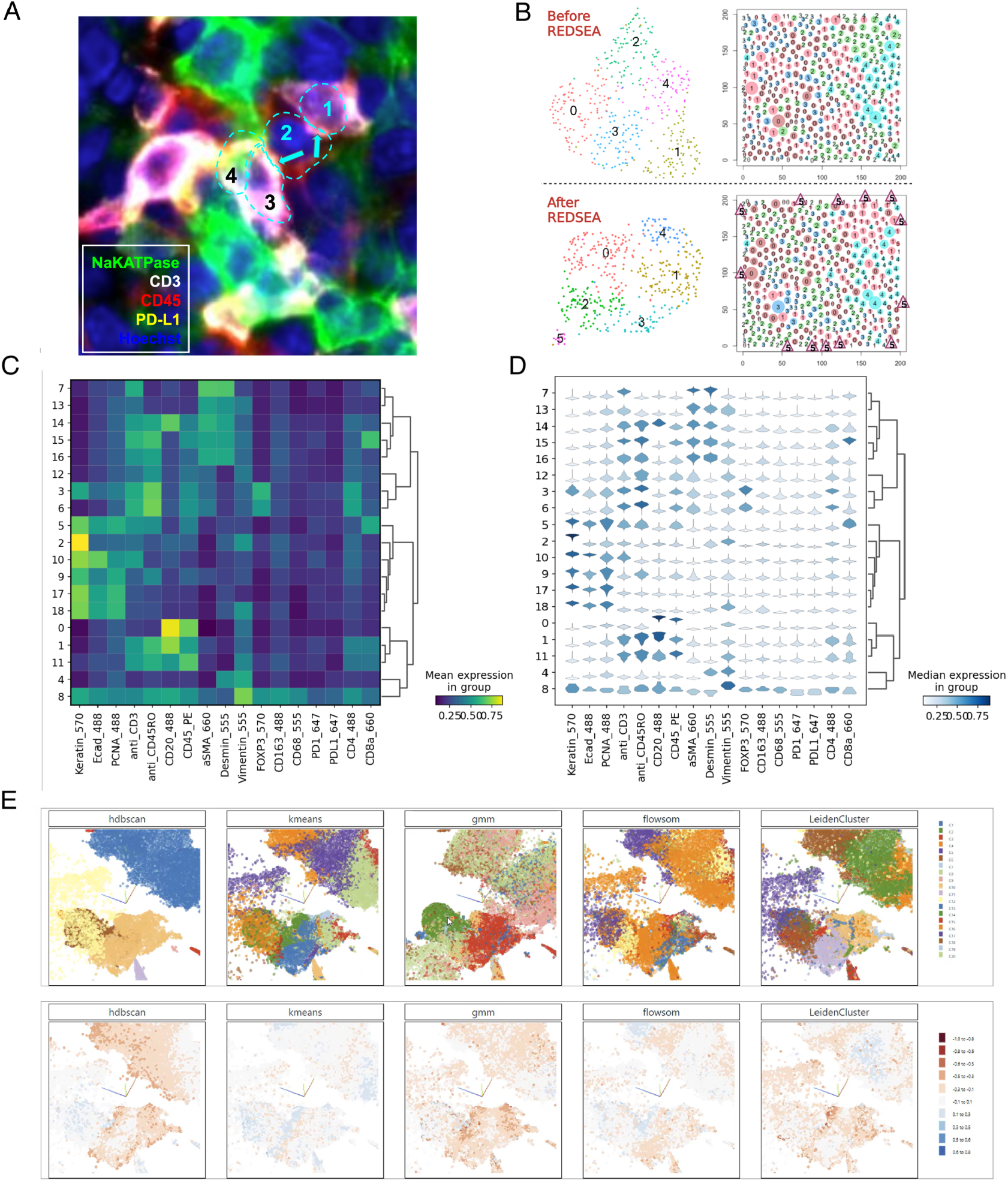
Spatial spillover and visual comparisons of cell type calling. **A**) Example of spatial cross-talk of adjacent cells in CyCIF stained images of tonsil. Boundaries of cells identified by segmentation are indicated by the dashed cyan lines and distinct cells are numbered. Pixel intensities from different markers are indicated by distinct colors. Spatial spillover of CD3 into adjacent cells is indicated by cyan arrows. **B**) Uniform Manifold Approximation and Projection (UMAP) of cell features and spatial representation of cells in a 200×200px tile before and after REDSEA. A novel Cluster 5 identified by REDSEA captures isolated cells at the image border (indicated by triangles). **C**) A traditional heatmap and (**D**) violin-matrix of cell data separated into clusters using the HDBSCAN^37^ algorithm. **E**) Visualizations generated by a web-based interactive tool for inspecting and comparing clustered data in a spatial context. Scatterplots of cells in UMAP embeddings with cells colored by cluster membership as a result of the respective clustering algorithms (*top row*) and colored by silhouette coefficients (*bottom row*). The plots are synchronized in navigation (zooming, panning, selections).

### Analysis of cell type classification

To visually and quantitatively assess cell type calling it is common practice to perform some type of clustering based on cell-specific features. However, the lack of ground truth and the diversity of possible preprocessing inaccuracies make clustering quality hard to judge using statistical summary measures. For effective quality control and intervention, it is thus essential to integrate biomedical researchers into the analysis loop. This can be achieved through data visualization and interactive interfaces, allowing experts to inspect and compare outcomes and to make decisions on which algorithm and parameter settings perform the best. A challenge to these visualizations is that there are numerous algorithms and data reduction techniques that can be used to group objects (cells) using similarity measures, each with specific advantages and disadvantages. Likewise, there are many approaches for comparing clusterings^38^, but none of them can be considered standard.

To overcome these limitations and facilitate comparisons of cell type clustering by multiple approaches, the challenge participants considered two alternative strategies: 1) static graphical representations of cluster quality, and 2) an interactive web-based platform for dynamic exploratory analyses.

The first approach involved creating a range of easily configurable summary heatmaps that provide more details on the clusters’ quality while maintaining a high-level overview of the entire dataset. Compared to traditional heatmaps (or matrices)^37^ that show mean marker values per cluster of cells (Figure 2C), violin-matrices additionally display marker distributions for each cluster (Figure 2D) as small multiples. Alternatively, marker intensity distribution within a cluster can be visualized in even more detail, on a single-cell level, by subdividing each cluster’s cells into thin stripes (Supplementary Figure 4). To gauge cluster quality even further, a color-encoding of computed silhouette scores enables rapid identification of clusters with, e.g., many cells of low silhouette score, a large variety of scores, or groups of low scoring instances within a cluster, indicating the cluster may be better split into smaller clusters.

The second approach involved the development of a fully interactive web application to explore multiple aspects of the data. The application allows inspection and simultaneous comparison of different clustering algorithms (k-means, density-based, Gaussian mixture models (GMMs), Self-Ordered Maps (SOMs), and Leiden) side-by-side in a spatial context, with coordinated zooming and panning of subregions within the plots, and coloring individual cells by cluster identity, expression value, or silhouette coefficients (Figure 2E). The web application scales to displaying large datasets with two or three UMAP axes and real-time adjustment of viewing angles and zoom ranges. This was achieved by utilizing scatter-gl^39^, a webgl-accelerated 2D and 3D scatter plot point renderer that is part of Tensorflow’s Embedding Projector^40^. The resulting tools are available on GitHub^41^ and can be installed on most personal computers.

### Summary, future work and remaining challenges to identifying and classifying cell types

Significant strides have been made in the development of robust image analysis tools that can effectively handle segmentation errors and image artifacts. However, further advancements are necessary to ensure their widespread effectiveness. One major challenge is the requirement for more training data with well-established ground truth to thoroughly evaluate and improve each implementation’s accuracy and generalizability. This is particularly true for the multilayer perceptron model, which was identified as the optimal solution for automatic detection of artifacts. With all deep learning models, additional training tends to improve their accuracy with the added costs of time and computational resources. An intriguing prospect lies in coupling automatic artifact detection with the ability to automatically correct these artifacts within the images. This integrated approach could provide substantial benefits, streamlining the analysis workflow and increasing overall accuracy.

To enhance the overall utility of each of these tools, it is crucial to improve their scalability and automation capabilities, as well as to evaluate their performance across diverse datasets encompassing various imaging modalities, resolutions, tissues, and segmentation methods to determine ideal conditions suited for each tool. Specifically toward addressing lateral spillover with REDSEA, the incorporation of non-membrane markers may reduce the reliance on manual parameter tuning.

Tools to rapidly evaluate and compare various clustering algorithms and the underlying feature data that gives rise to the different resulting clusters are needed. This is true not only for imaging data but for high-dimensional clustering, in general, including single-cell RNA sequencing data^42–45^. Silhouette-heatmaps offer one solution for static visualization, but other metrics and visualizations may also be considered. Additionally, different cluster quality measures could be incorporated into the web-based visualization tool to aid in assessment of clustering outcomes.

### Image representation learning

Emerging multiplexed imaging technologies create images consisting of a large number of markers and provide high-dimensional single-cell feature tables, but analyzing single-cell multiplex imaging data is still primarily limited to extracting a single mean intensity value per channel, per cell. This classical image feature extraction approach is biased toward known and easily measured features, does not fully leverage multiplex imaging information, and can miss subtle but important subcellular features, such as marker polarity and staining colocalizations across markers that might indicate divergent cell states. Improving the number, nuance, and kinds of features extracted from multiplex imaging will improve phenotyping and give researchers a better understanding of cell states, cell population heterogeneity, cell-cell communication, and intercellular regions. However, the financial and temporal costs of multiplexed imaging approaches are higher compared to the traditional imaging modalities, such as hematoxylin and eosin (H&E) staining and brightfield microscopy. To bridge the gap, recently-proposed deep learning approaches can extract relevant morphological features from low-dimensional (e.g., H&E stained) images that are not easily perceived by the human eye and use those features to infer the expression of molecular markers; these approaches are collectively referred to as virtual staining^46–48^. It remains a challenge to provide such data for human visualization in a convenient and browsable manner. Although methods for photography, brightfield imaging, and low-plex immunofluorescence data generate RGB thumbnails by downsizing and cropping, these are not immediately generalizable for data with 100 channels without losing interpretability. Here we discuss methods for feature extraction using variational autoencoders (VAEs), automated summarizing of high dimensional data with thumbnail images, and predicting marker staining intensities from H&E images.

### Feature extraction with VAEs

VAEs have previously been successfully trained on various biomedical data modalities such as bulk and single-cell gene expression, and imaging data^49^. A potential challenge to VAE feature extraction from single-cell imaging data is an abundance of unimportant or uninformative morphological features driving differences between biologically similar images and skewing the results in undesired ways^11, 50^.

Hackathon participants trained several forms of variational autoencoders (VAEs), including standard VAE, β-VAE^51^, invariant conditional VAE (C-VAE)^52^, and Multi-Encoder VAE (ME-VAE)^11^, on immunofluorescence images of human mammary MCF10A cells treated with TGF-β or PBS (control) and evaluated their ability to separate biologically distinct cell populations based on latent features as well as standard morphological and spatial features.

The participants found that most VAEs were confounded by uninformative features, with the exception of ME-VAE, which showed strong discriminatory performance in an unsupervised setting, improved further by subsetting the latent space to the top 10 most highly variable features (Figure 3A).

**Figure 3:**
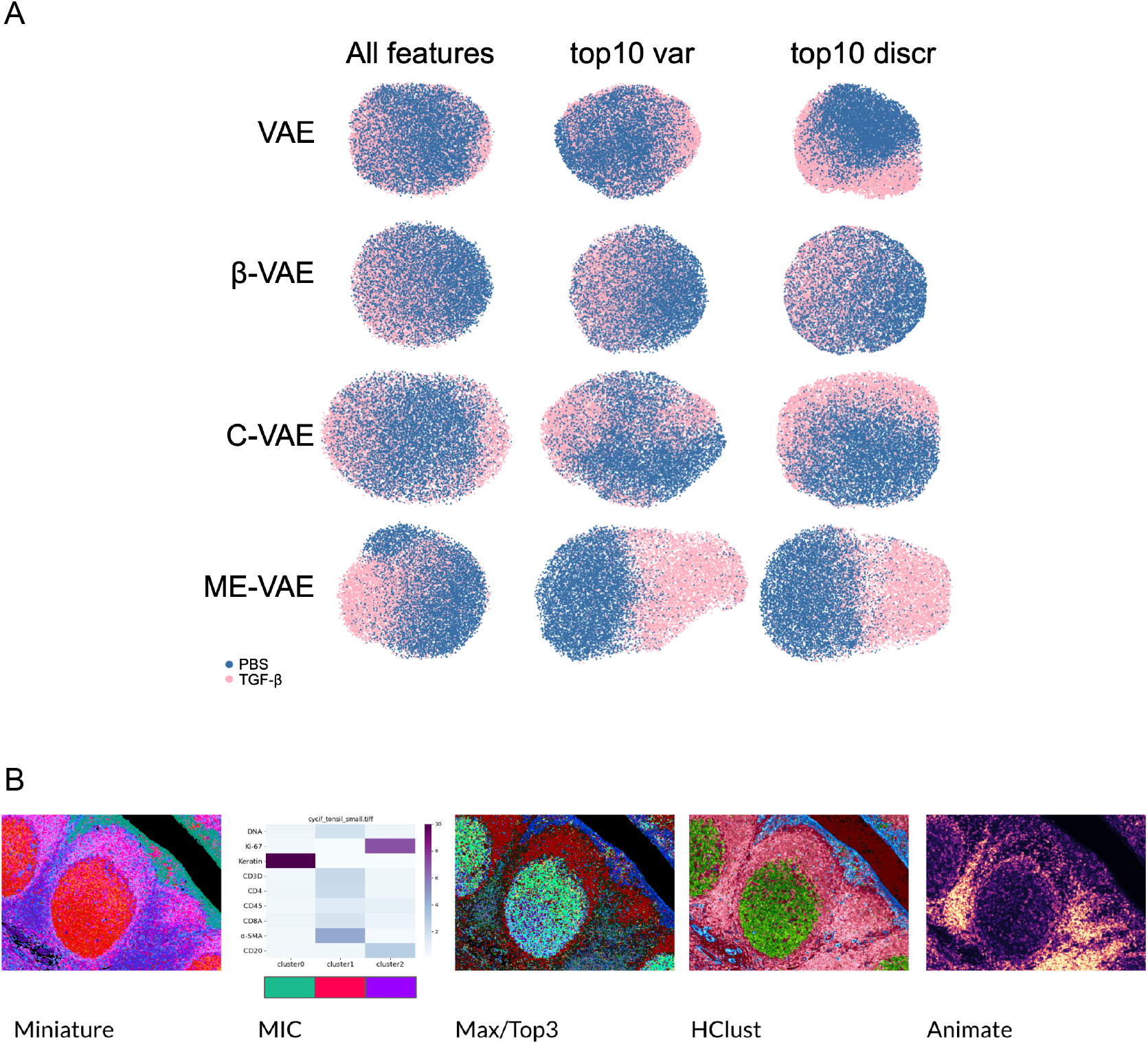
Image representation learning by VAEs and for thumbnail generation. **A)**. Each implementation of VAE was qualitatively assessed for their ability to distinguish control-(PBS-)treated from TGF-β-treated MCF10A cells using all morpho-spatial features or the top 10 variable (var) features compared to preselecting the top 10 discriminatory (discr) features extracted from the images. Feature space is reduced to two dimensions using UMAP embedding. Class labels of TGF-β- or PBS-treated cells are shown in pink and blue, respectively. **B)** Example thumbnail images. Each panel shows a thumbnail (or associated comparative plot) generated by the methods described in the main text (panel labels). All approaches were applied to a 0.9 mm^2^ (9 megapixels) 9-channel CyCIF image of a human tonsil germinal center.

### Thumbnail generation

Thumbnails are scaled down representations of larger images that enable easier viewing, and faster access, storage and management. Since these are miniature versions of large images, challenges arise in techniques related to downscaling, compressing, and resizing these larger images, while taking into account both noise as well as the higher number of the multiplex channels.

Four different methods were developed to compress highly multiplexed images into three-color (e.g., RGB) thumbnails to facilitate interpretation. These made use of two pre-existing tools as starting points: Auto-Minerva^53^, which uses a GMM to isolate tissue foreground signal, and Miniature^54^, which employs UMAP dimensionality reduction on image pixels to embed images in the CIE L*C*h color space while ensuring image regions with similar protein expression patterns receive similar colors. The methods were evaluated on three CyCIF datasets that included two whole-slide images of colorectal cancer and healthy tonsil, and one tissue microarray^10^, which was acquired at 0.65 microns per pixel with a 20X/0.75NA objective lens.

The first method, called MIC (Figure 3B), builds on Miniature^54^ by performing k-means clustering on the UMAP embedding, followed by calculating a variable importance metric, which can be displayed in a compact heatmap, enabling rapid assessment of markers driving heterogeneity within an image; the heatmap is complementary to a thumbnail and aids its interpretation. In the tonsil CyCIF dataset, it was seen that the three clusters were driven primarily by Keratin, α-SMA and Ki-67. The next method, Max/Top3, explored a maximum intensity projection approach taking the top three most highly-expressed channels at each pixel and assigning them to the red, green, and blue channels of the thumbnail. HClust is a hierarchical clustering approach to find three informative channel groups per image, within which intensities are aggregated and colored in the CIE L*C*h colorspace. Several aggregation and color assignment methods were explored including the group hue, pixel maximum and pixel sum, in combination with the cut point of the dendrogram. Finally, Animate generates an animation that cycles through all channels after downsizing and auto-thresholding.

### Virtual immunofluorescence staining

Although multiplex immunofluorescence staining can provide deep insights into quantitative and spatial aspects of cancerous tissues, it can be prohibitively expensive and time consuming, and repetitive processing steps risk introducing artifacts, such as tissue degradation during later cycles. Alternatively, it may be possible to infer the expression of specific protein markers from tissue autofluorescence using image-to-image translation, i.e., virtual staining. Ideally, by leveraging the endogenous autofluorescence of human tissues, these techniques can recreate pathology images without requiring arduous chemical staining procedures^48, 55^ and can therefore expand the utility of digital pathology. An extension of this approach is the prediction of specific marker expression from unstained images—virtual IF^46, 48^.

The objectives for the hackathon were to use the SHIFT implementation^56^ of the pix2pix^57^ conditional generative adversarial network model to translate label-free autofluorescence to immunofluorescence signals. The training data were whole-slide images of healthy human kidney tissue samples with three channels of autofluorescence images and 10 different immunofluorescence stains of kidney-specific markers, acquired by serial sectioning [verify]. Training SHIFT to predict the proximal tubule marker AQP1 from the label-free autofluorescence images produced a virtually stained image that, when analyzed for single-cell expression of AQP1, had correlated levels of predicted expression (Pearson ρ = 0.48; Supplementary Video 1) to those measured on the neighboring section stained for AQP1. However, the low autofluorescence detected in the nuclei was insufficient for SHIFT to predict nuclear staining.

### Summary, future work and open questions of image representation learning

VAEs are powerful tools for extracting and representing latent variables from high-dimensional images and enabling interpretation of the learned representations. However, applying VAEs to extract biologically meaningful information from single-cell imaging is frequently driven by unimportant or uninformative features. By using a multiple-encoder VAE and focusing on the top ten most variable features of images, hackathon participants were able to substantially increase the discriminatory power of VAEs. While methods like ME-VAE can learn representations without explicit labels, more effective unsupervised learning algorithms that can discover informative features without relying on prior knowledge remain an open challenge. Representation learning methods that can capture meaningful and transferable features that generalize well across different tissue types, diseases, and multiplex imaging platforms may be useful in this regard. Ideally, the learned representations will be biologically interpretable and meaningful, which would add trust in these machine learning approaches. The development of representation learning methods that produce interpretable and explainable features is an intensely active area of research and progress along this front should provide powerful new tools for cancer image analysis.

Although several methods of thumbnail generation and channel-to-color associations were developed, their suitability across applications will differ. These applications may include interactive data analysis and visualization (such as Mistic^58^), online data portals [such as those belonging to HTAN, HuBMAP, and NCI’s Imaging Data Commons (IDC)], and local file browsers. However, any use case is likely to have a unique feature set that would require prioritizing one thumbnail generation method over another. Thus, thorough testing will be required to assess whether specific tasks can be completed with more ease and accuracy.

The automated translation of one domain (e.g., autofluorescence images) to another (e.g., immunofluorescence images) is not yet easily implemented, especially when compared to deep learning approaches for segmentation. Fine tuning of models remains critical to their successful implementation. However, success remains highly subjective, based primarily on visual inspection. An alternative approach that would provide quantitative evidence to support “success” would be to compare the outcome of downstream analyses, like cell segmentation, between a virtual and true immunofluorescence images. Many downstream analyses that currently rely on true fluorescence images could act as useful quantitative metrics for determining the accuracy of virtual staining.

### Image processing at scale

Modern highly-multiplexed imaging methods are capable of producing TB-scale datasets, with individual images requiring dozens of GB of storage and extensive resources for processing whole-slide images in an end-to-end pipeline. The scale of today’s images poses substantial challenges for applying existing methods originally designed on smaller datasets. Hackathon challenges designed to address this theme included optimization of existing methods to increase throughput efficiency, end-to-end image analysis with Galaxy^59–61^, and importing tissue volumes into Neuroglancer^62^, an open source volume visualization tool developed by Google.

### Deploying image segmentation at scale

The effective application of image analysis at scale requires a multi-pronged approach with advancements in three areas: algorithm optimization, identification and exploitation of parallelism, and scalable deployment on cloud infrastructure. The need for interactive visualization and human-in-the-loop workflows complicates all three areas. This stands in contrast to other molecular data modalities, such as RNA sequencing, where operations like sequence alignment and variant calling can be carried out without continuous integration with data visualization and user feedback^63, 64^.

Hackathon participants examined CellPose^9^, a popular neural-net-based cell segmentation method, for ways in which its processing efficiency could be improved for applications to large datasets. Because localization of cells in images is a central step in any cancer image analysis pipeline, inefficient cell segmentation methods can present a substantial bottleneck. The participants first profiled individual aspects of the method (file I/O, neural net inference, and post-processing of model outputs) to determine computational bottlenecks. The participants also measured how the run time scales with image size and attempted to identify opportunities for parallelization. All analyses were carried out using a CyCIF tissue microarray dataset^10^. Participants determined that file I/O was not a significant bottleneck and that over ⅔ of runtime was devoted to neural network inference, which scaled linearly with image size within the limits of available memory. GPU-based computation within CellPose resulted in a 17.6-fold improvement in runtime compared to CPU-based processing. A further four-fold reduction in runtime was achieved by removing Cellpose’s redundant models that were all trained identically but initialized from a different set of random weights. Removing these was not detrimental to segmentation accuracy, which aligns with the general view that an effective ensemble should contain weak, complementary predictors^65^.

### End-to-end image analysis with Galaxy

Processing image datasets using scalable and standardized workflows enables reproducible analysis and provides harmonized and comparable results across imaging modalities. Galaxy is a web-based platform widely used for large-scale data processing and analysis^59–61^ and is compatible with multiple computing infrastructures, which allows it to leverage cloud computing resources and Unix/Linux based high performance computer clusters to power large-scale data analysis. By coupling image processing, analysis, and visualization, the entire workflow can be executed remotely with no need for downloading large files locally or moving data to external software. The Galaxy-MTI tool suite has recently been integrated with the core MCMICRO^10^ pipeline to enable automated and reproducible image processing and analysis but has not yet been broadly deployed. The objectives of hackathon participants were to expand the Galaxy/MCMICRO workflow to function with imaging modalities beyond CyCIF. This included preprocessing and registration of multiplexed immunohistochemistry (mIHC) images using PALOM^66, 67^ and comparing the segmentation output of two different algorithms (StarDist^68, 69^ and Ilastik^70^) within Galaxy.

CyCIF datasets from a breast tissue microarray and tonsil, both sampled at 0.65 micrometers/pixel, were used as reference images on which the various processing steps were performed. The hackathon participants developed several pipelines and extensions to the software suite, including adding several image preprocessing/registration and segmentation algorithms, developing a new workflow optimized for non-fluorescence based multiplex tissue imaging platforms, integrating PALOM for piecewise alignment of images^66, 67^, and attempting to deploy StarDist^68, 69^ for 2D segmentation of CyCIF images. (StarDist was mostly implemented but resulted in errors as different steps in the pipeline that require further development.) To facilitate portability, all tools were built within Docker images.

### Scalable Visualization of 3D data using Neuroglancer

The challenge objectives were to import, process, render, and navigate multichannel 3D volumetric datasets in Neuroglancer^62, 71^. Participants were given 3D datasets of highly multiplexed CyCIF^8^ melanoma datasets, each spanning 500 × 500 × 5 micrometers, from the Pre-Cancer Atlas. Each region was sampled at 108 nm lateral and 200 nm axial resolution. These samples displayed a wide array of histopathological features associated with early stage melanomas including brisk immune infiltrates, tumor regression, and pagetoid spread.

Participants converted each channel in the data into a separate 3D segmentation mask using global thresholding or machine-learning-based pixel classification methods. From here, the segmentation masks were converted to meshes using marching cubes^62, 72^. The meshes could then be loaded and displayed in Neuroglancer. To simultaneously look at multiple channels, participants loaded multiple meshes into a single view and overlaid them with the ability to enable or disable individual channels. In addition to 3D surface rendering, participants used a 2D slice view to render a specific slice of the multi-volume selectable by scrolling through the third axis. The 2D view shows the precise image data, but allows overlaying the segmentation mask in semi-transparent layers (**Figure 4**). Both the 2D and 3D views can be interactively linked for synchronous navigation.

**Figure 4.**
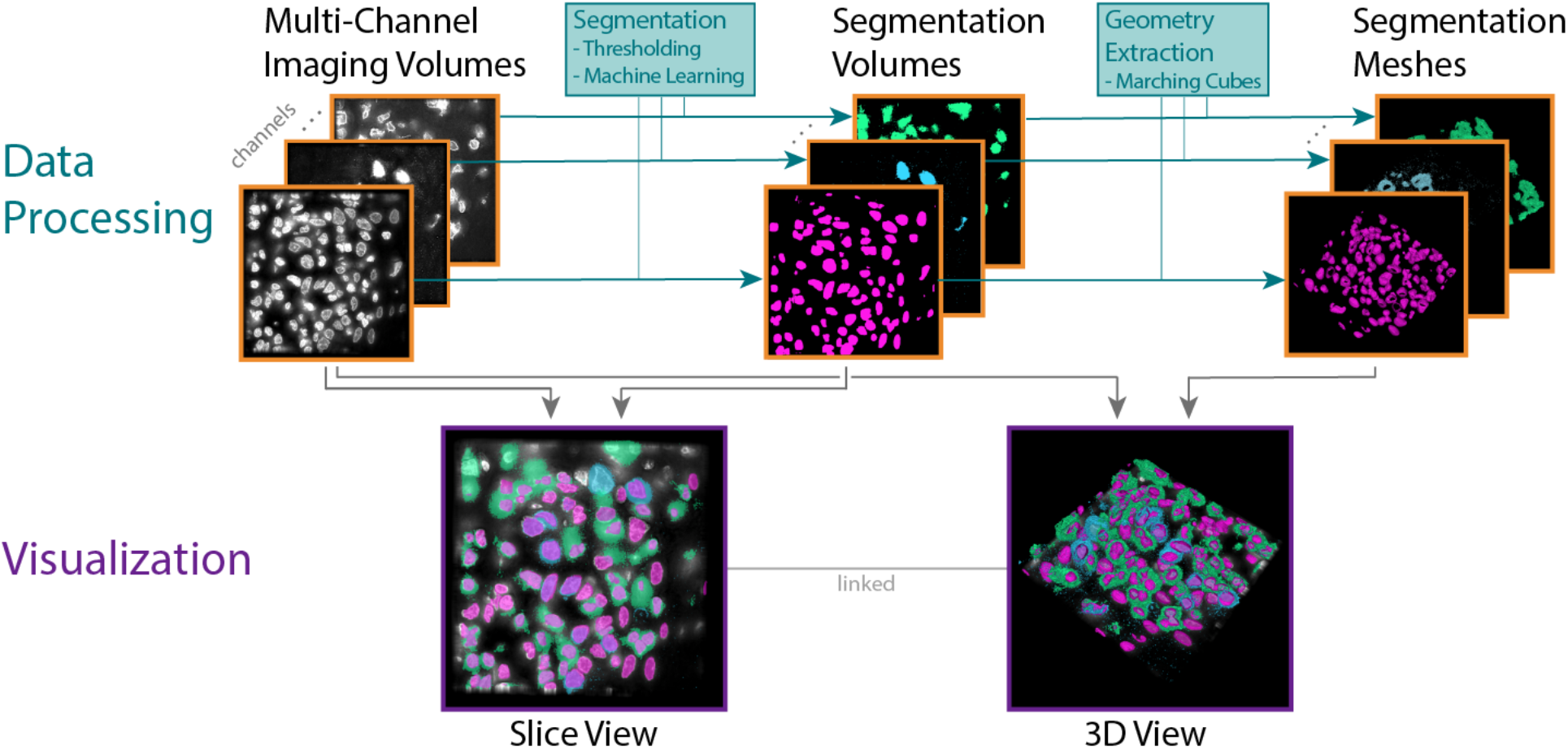
Data Processing and Visualization pipeline developed during the challenge for Neuroglancer. Highly multiplexed CyCIF data are stored as multi-channel imaging volumes (*top, left*), where each volume represents one channel. For simplicity, volumes are depicted as single slices in this figure. Each volume is segmented, either via thresholding or more complex machine learning approaches and stored as binary segmentation volume (*top, middle*). Subsequently, for each segmentation volume (i.e., segmented channel) the geometry of the segmented structures is extracted and stored as a geometry mesh for subsequent 3D surface rendering (*top, right*). The visualization pipeline supports a slice view that can combine an original imaging volume with several segmentation volumes (*bottom, left*) and a 3D view (*bottom, right*). The 3D view can represent the volume as extracted meshes or a clipping plane.

### Summary, future work and open questions

The incorporation of new tools in Galaxy is relatively straightforward due to its highly modular architecture and the use of Docker containers. However, the generation of Docker containers that are fully compatible with the Galaxy system and enable full functionality of the embedded resources required diligence. Although progress was made along multiple fronts to develop new Galaxy pipelines, none of them have been thoroughly tested and hardened. Future efforts will specifically focus on enabling users of Galaxy to apply the new implementations to various data sets to ensure their proper functionality and on training new neural net (or other deep learning) models for segmentation tasks that can be deployed more broadly.

Many of the tools and platforms are optimized for fluorescence based microscopy platforms, but there is a growing need to expand software tools to flexibly handle multiple imaging modalities. Additional enhancements may include more sophisticated image registration and image pre-processing methods, robust testing, and more efficient resource allocation. Neuroglancer was found to be suitable for scaling up multichannel 3D datasets. However, it is still dependent on provision of high quality 3D segmentation masks especially where there are highly intricate biological structures. To make Neuroglancer more computationally efficient, one could adopt the more computationally efficient webGPU API and use more scalable data formats like Neuroglancer Precomputed^73^ that negate the need of loading the entire image volume into working memory.

### Overall Summary

The preparation for the IAWG hackathon involved a critical review and summary of the many remaining challenges to automating image processing and analysis for digital pathology, some of which we attempted to specifically address within the hackathon. While the grouping of the challenges into themes was primarily out of convenience, the three main categories appear particularly relevant to most approaches that attempt to extract quantitative single-cell measurements from the numerous techniques that have been developed to generate high-dimensional multiplexed images of cancerous tissues. The first theme addresses challenges to cell type classification and assessment and includes the detection and removal of technical artifacts from images and downstream analyses, and how to visually and quantitatively assess cell classifications. Since it is impossible to prevent all artifacts from entering into the process, it is important to acknowledge their potential for introducing errors into downstream analyses. Having tools to automatically and accurately identify these potential sources of error will enable the quantification of their potential contribution to errors in interpretation and ideally make their effects negligible.

The second theme related to translation and visual representation of spatial aspects of high dimensional imaging data. While the approaches considered during the hackathon focused on single-cell variability, their further expansion to cell neighborhood and tissue-level spatial information, as well as integration with other data modalities, will be of high importance to a better understanding of cancer. The techniques described here generally focus on translating or reducing the information from a particular modality into another domain with the primary objectives of identifying and understanding the relationships between the domains and to facilitate their visual interpretation. We expect standardizing and hardening these approaches will enhance our ability to interpret large datasets in an automated manner.

The last theme dealt with the challenges of big data and the scaling of digital image analyses to accommodate them. Prior focus on some steps in the image processing and analysis pipeline, such as cell segmentation using CellPose^9^, appears to have succeeded in scaling well with increasing data. However, the quality and accuracy of cell segmentation still requires substantial optimization for different datasets, which dramatically reduces the computational efficiency gains. Resources like Galaxy^60^ can assist with the process of optimizing the steps by providing a platform for executing reproducible pipelines, including quality control tests, using a multitude of computational tools in parallel. We strongly support the use of tools that enhance reproducibility and enable cross compatibility, especially with the abundance of different platforms and data types associated with cancer tissue imaging.

The virtual hackathon held in 2022 provided our community with the ability to make progress toward multiple challenges to fully automating the analysis of highly multiplexed tissue data—a lofty and possibly unattainable goal. However, there still remain additional open questions and areas of research that will only be amplified by the advancement of high-throughput multiplexed imaging, computational methods, and machine learning architectures. Some of these challenges include: how best to integrate spatial imaging data from fixed tissues with other single-cell modalities such as single-cell sequencing data; how to analyze and provide broader access to large 3D imaging datasets; how to interpret results from spatial and neighborhood analyses, such as those applied to studying tumor-immune microenvironment interactions; and how to link the static fixed tissue imaging data to our understanding of the highly dynamic evolutionary process that is cancer.

## Acknowledgments

The hackathon was generously supported by sponsors Carl Zeiss, Lunaphore Technologies, and MARK III Systems through prizes for hackathon winners. The authors, hackathon organizers, and participants are funded by NIH grants 3U54CA225088-04S1, U2C-CA233262, U54CA225088, U54CA209988, U2CCA233280, U54CA217450, R50CA243783, U54CA217450, NSF grant NCS-FO-2124179, NSF grant IIS-1901030. We would also like to acknowledge the excellent work performed by several of the participants who were also awarded prizes for their efforts during the hackathon, including Edward Novikov, Laurent Bataille, and Monica Del Valle. We would also like to acknowledge the contribution of several participants with whom we have lost contact and have been unable to include them as authors on the manuscript, including: Luke Sargent, Laurent Bataille, Walid Bousselham, Behnaz Bozorgui, Elmar Bucher, Chamika Gangul, Qiang Gu, and Jason Lu.

## SUPPLEMENTARY INFORMATION

Image data used for each challenge addressed during the hackathon are available at https://www.synapse.org/#!Synapse:syn26848022/files/ A list of all acronyms and abbreviations and a list of all URLs to relevant code (GitHub) repositories are provided as Supplementary Text.

