## Supplementary Text for "Addressing persistent challenges in digital image analysis of cancerous tissues"

### Code repositories

#### Challenges to identifying and classifying cell types

Automatic artifact detection from the spatial feature table:
https://github.com/IAWG-CSBC-PSON/hack2022-01-artifacts

Artifact correction/suppression directly on images:
https://github.com/IAWG-CSBC-PSON/hack2022-11-cosmetic

Lateral spillover correction using REDSEA:
https://github.com/IAWG-CSBC-PSON/hack2022-05-cross-talk

Analysis of cell type classification:
https://github.com/IAWG-CSBC-PSON/hack2022-06-viz-comp

#### Image representation learning

Feature extraction with VAEs:
https://github.com/IAWG-CSBC-PSON/hack2022-04-vae

Thumbnail generation:
https://github.com/IAWG-CSBC-PSON/hack2022-08-thumbnails

Virtual immunofluorescence staining:
https://github.com/IAWG-CSBC-PSON/hack2022-03-virtual-if

#### Image processing at scale

Deploying image segmentation at scale:
https://github.com/IAWG-CSBC-PSON/hack2022-07-tb-scale

End-to-end image analysis with Galaxy:
https://github.com/IAWG-CSBC-PSON/hack2022-09-galaxy

Scalable Visualization of 3D data using Neuroglancer:
https://github.com/IAWG-CSBC-PSON/hack2022-10-neuroglancer

**List of Abbreviations and acronyms**

2D: Two-dimensional
3D: Three-dimensional
4i: iterative indirect immunofluorescence imaging
α-SMA: Alpha Smooth Muscle Actin
API: Application Programming Interface
ARTNet: Acquired Resistance to Therapy Network
CCBIR: Cellular Cancer Biology Imaging Research
CD3: Cluster of differentiation 3
CIE L*C*h: CIELCh color scale
CODEX: Co-detection by indexing
CPU: Central Processing Unit
CSBC: Cancer Systems Biology Consortium
C-VAE: Conditional Variational Autoencoder
CyCIF: Cyclic immunofluorescence
DFR: Deep Feature Reconstruction
EMIT: Exemplar Microscopy Images of Tissues
GAN: generative adversarial networks
GB: Gigabyte
GMM: Gaussian Mixture Model
GPU: GraphicsProcessing Unit
H&E: hematoxylin and eosin
HCA: Human Cell Atlas
HDBSCAN: Hierarchical Density-Based Spatial Clustering of Applications with Noise
HTAN: Human Tumor Atlas Network
HuBMAP: Human BioMolecular Atlas Project
IAWG: Image Analysis Working Group
IDC: Imaging Data Commons (from NCI)
IF: ImmunoFluorescence
IMC: Imaging Mass Cytometry
iMS: imaging mass spectrometry
I/O: Input/Output
Ki-67: Marker Of Proliferation
K-means: A standard data clustering method where the number of clusters is denoted by ‘K’
LDA: Linear discriminant analysis
LightGBM: Light Gradient Boosting Machine
MCF10A: Michigan Cancer Foundation mammary epithelial cell line
MCMICRO: Multiple-choice microscopy pipeline
MERFISH: Multiplexed Error-Robust Fluorescence In Situ Hybridization
ME-VAE: Multi-Encoder VAE
mIHC: multiplex immunohistochemistry
MLP: Multi-Layer Perceptron
MSE: mean squared error
MTI: multiplex tissue imaging
mIF: multiplexed immunofluorescence
MxIF: multiplexed immunofluorescence
NCI: National Cancer Institute
PALOM: Piecewise alignment for layers of mosaics

PBS: phosphate-buffered saline
PCA: principal component analysis
PDXNet: PDX (patient-derived xenografts) Development and Trial Centers Research Network
PLS-DA: partial least squares-discriminant analysis
PR: precision
PSNR: peak-signal-to-noise-ratio
PS–ON: Physical Sciences–Oncology Network
QC: quality control
QDA: quadratic discriminant analysis
RC: recall
REDSEA: REinforcement Dynamic Spillover EliminAtion
RF: Random Forest
ROC: Receiver operator curve
seqFISH: Sequential Fluorescence In Situ Hybridization
SHIFT: speedy histological-to-immunofluorescent translation
SOM: Self-organizing Map
StarDist: cell segmentation method using Star-convex Polygons
SVM: support vector machines
TB: Terabytes
TGF-β: Transforming growth factor beta
TREAT-AD: TaRget Enablement to Accelerate Therapy Development for Alzheimer's disease
XGBoost: eXtreme Gradient Boosting
UMAP: Uniform Manifold Approximation and Projection
VAE: variational autoencoder
