## Supplementary Figure 1 for "Addressing persistent challenges in digital image analysis of cancerous tissues"

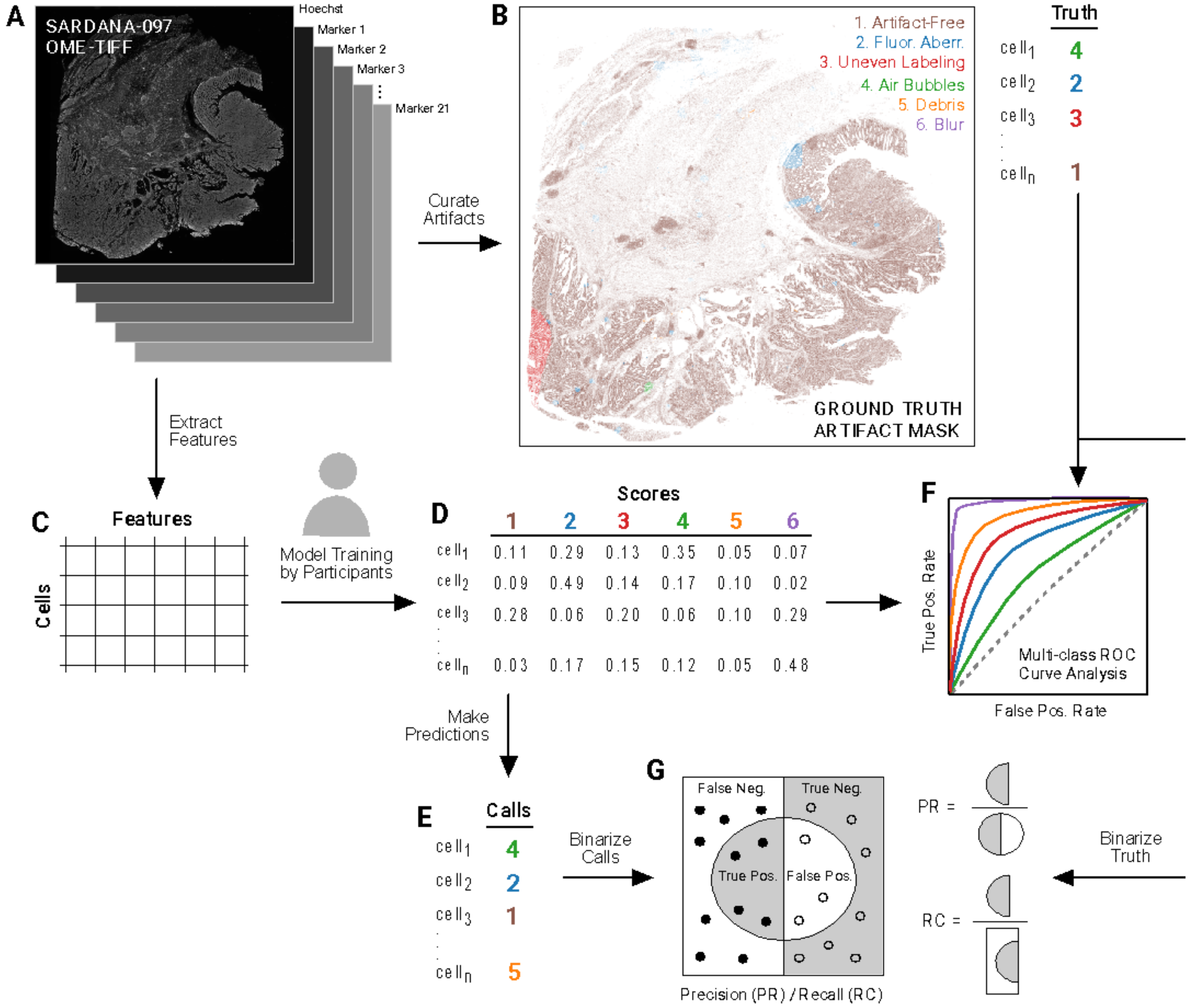


**Supplementary Figure 1. Overview of approach to address challenges to automatic artifact detection**. (**A**) Schematic representation of the multi-channel colorectal serial sectioned CyCIF dataset used for this challenge showing the Hoechst channel. (**B**) Ground truth artifact mark (left) and corresponding truth table (right) for artifacts in the 97th serial section of the colorectal CyCIF dataset colored by whether cells are artifact-free (brown) or are affected by 1 of 5 classes of visual artifacts: fluorescent contaminants (blue), uneven antibody labeling (red), coverslip air bubbles (green), slide debris (orange), image blur (purple). (**C**) Schematic representation of single-cell features derived from the colorectal CyCIF dataset. (**D**) Classifier probabilities for cells in the colorectal CyCIF dataset. (**E**) Classifier prediction calls derived from classifier scores tables (panel D). (**F**) Multi-class ROC analysis for each of the 5 artifact classes in the colorectal CyCIF dataset. (**G**) Precision and recall computations for individual and combined artifact classes.
