## Supplementary Figure 2 for "Addressing persistent challenges in digital image analysis of cancerous tissues"

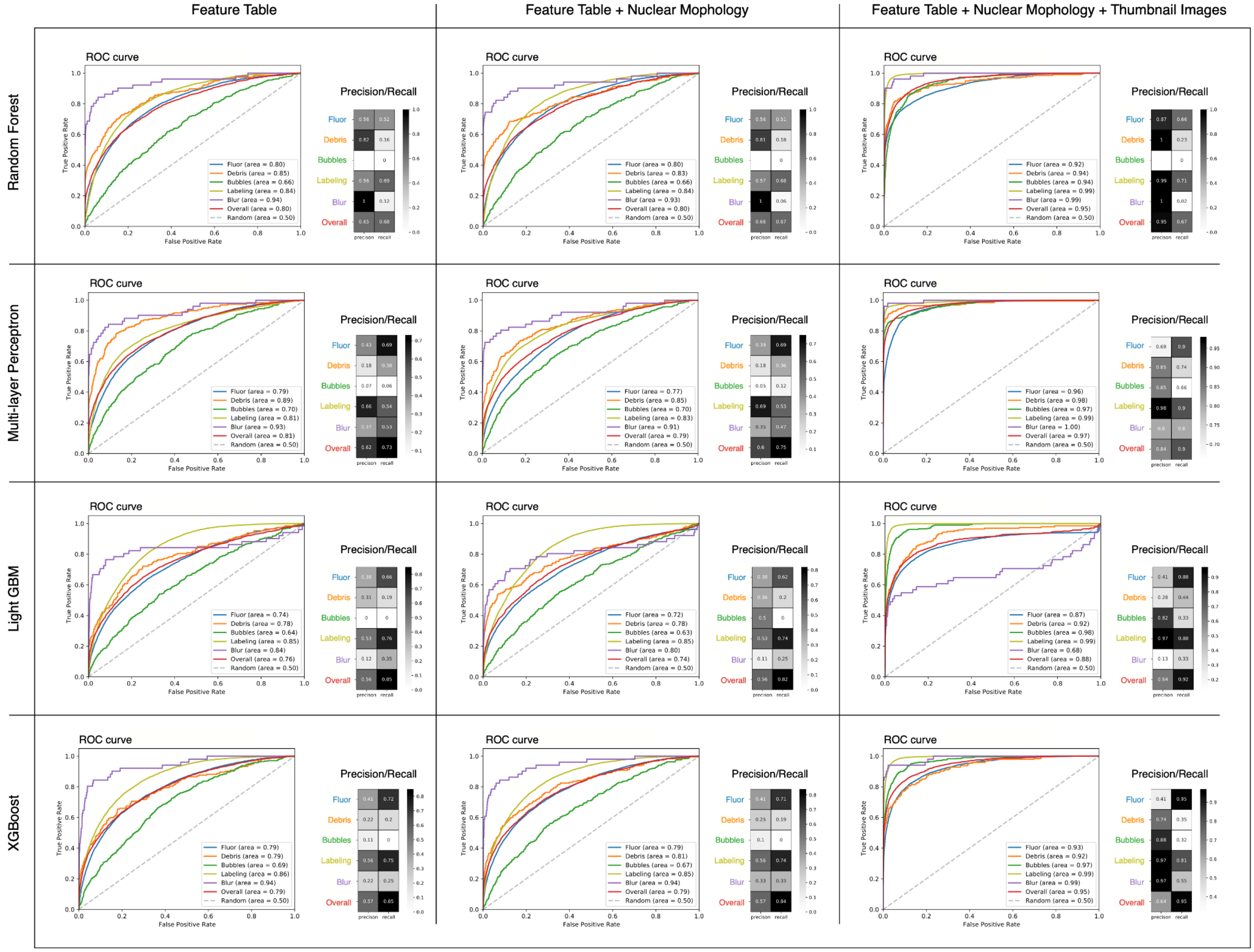


**Supplementary Figure 2**. **Artifact Classifier Performance**. Multi-class ROC curve analysis (line plots), and metrics of precision (PR) and recall (RC) shown as heatmaps for 4 probabilistic machine learning classifiers (rows) trained on 3 different feature sets (columns). These feature sets are referred to as FS1, FS2 and FS3 in the main text.
