## Supplementary Figure 3 for "Addressing persistent challenges in digital image analysis of cancerous tissues"

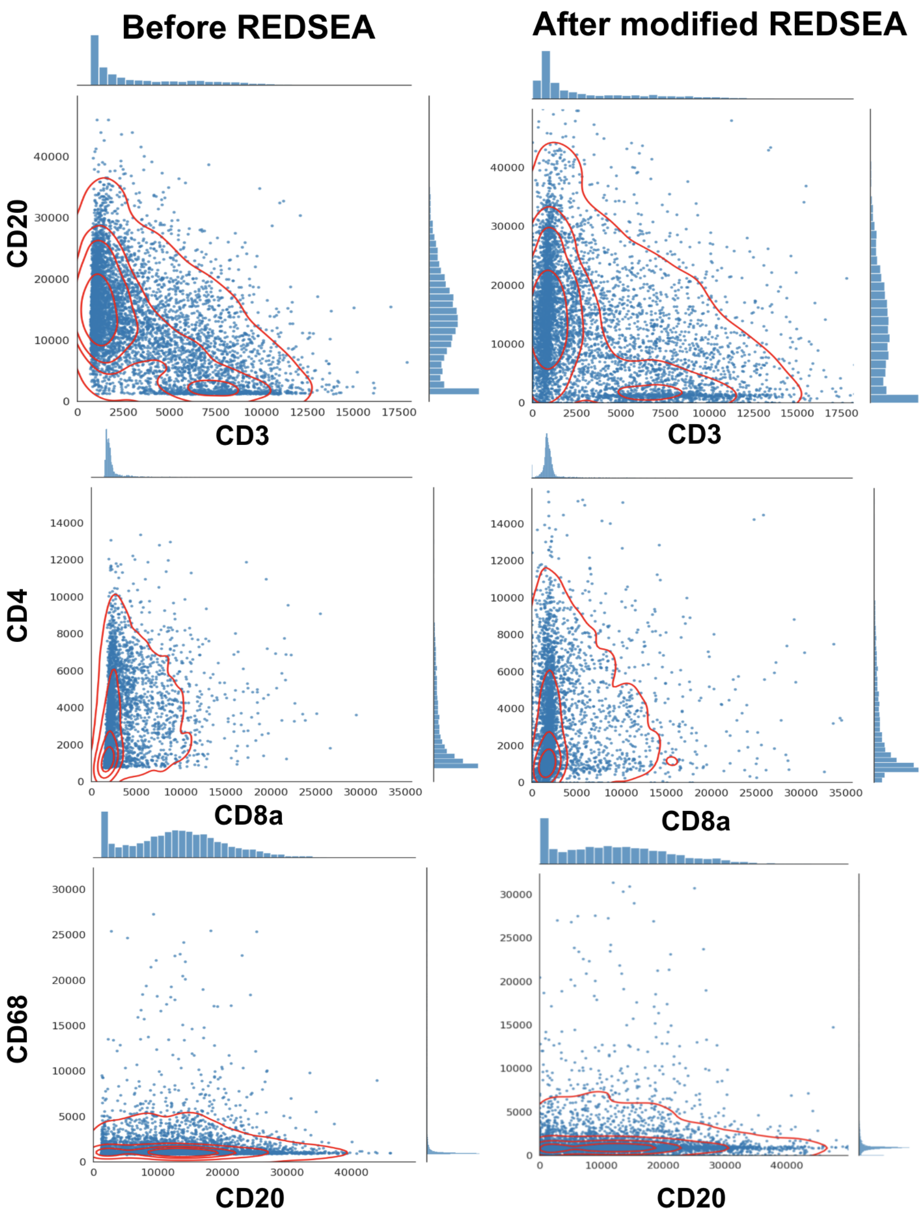


**Supplementary Figure 3: Comparison of cell data before and after application of REDSEA.** Co-expression plots for a set of mutually exclusive markers (CD20 vs anti-CD3, CD4 vs CD8a, CD68 vs CD20). Cells are represented by blue points (n=6390). Isodensity contours for cells are traced by the red lines. The percentage values for double-positive and single-positive cells were not calculated because the thresholds used in the original REDSEA are unknown. However, the density histograms (placed above and right for each co-expression plot) indicate that the proportions of single-positive cells were maintained or increased by the modified implementation of REDSEA.
