## Supplementary Figure 4 for "Addressing persistent challenges in digital image analysis of cancerous tissues"

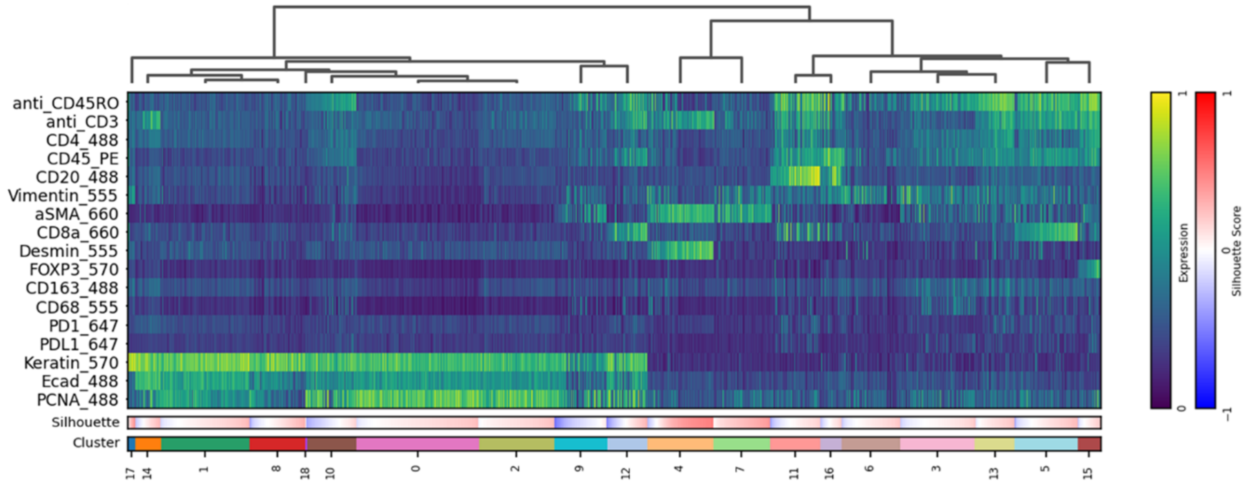


**Supplementary Figure 4: Detailed heatmap for marker expressions at single-cell resolution with color bars representing silhouette coefficients and cluster memberships**. Marker expression levels and silhouette scores are represented by colors shown in scale bars on the right.
